# Membrane-tethered peptides derived from intracellular loops 2 and 3 of the urotensin II receptor act as allosteric biased ligands

**DOI:** 10.1101/2020.11.23.394270

**Authors:** Hassan Nassour, Tuan Anh Hoang, Ryan D. Martin, Juliana C. C. Dallagnol, Étienne Billard, Myriam Létourneau, Ettore Novellino, Alfonso Carotenuto, Bruce G. Allen, Jason C. Tanny, Alain Fournier, Terence E. Hébert, David Chatenet

**Affiliations:** Institut National de la Recherche Scientifique, Centre Armand-Frappier Santé Biotechnologie, Groupe de Recherche en Ingénierie des Peptides et en Pharmacothérapie (GRIPP), Université du Québec, Ville de Laval, Québec, Canada; Department of Pharmacology and Therapeutics, McGill University, Montréal, Québec, Canada; Department of Pharmacy, University of Naples “Federico II”, Naples, Italy; Department of Medicine, University of Montreal, Montreal Heart Institute

**Keywords:** G protein-coupled receptors, pepducins, lipidated peptides, urotensin II receptor, allosteric modulators, cellular signaling

## Abstract

Over the last decade, the urotensinergic system has garnered significant attention as a promising new target for the treatment of various cardiovascular diseases and also cancer. Significant investment toward the development of clinically relevant UT ligands for therapeutic intervention has been made but have met little to no success to date. The UT system, which has yet to be effectively targeted, therefore remains to be therapeutically exploited. The discovery of allosteric sites that allow modulation of receptor activity will increase the searchable chemical space against a disease-relevant target. Pepducins and other lipidated peptides have been used as both mechanistic probes and potential therapeutics. Therefore, pepducins derived from the human urotensin II receptor might represent unique tools to generate signaling bias and study UT signaling networks. Two hUT-derived pepducins, derived from the second and the third intracellular loop of UT, respectively, have been synthesized and pharmacologically characterized. Our results demonstrated that hUT-Pep2 and [Trp^1^, Leu^2^]hUT-Pep3 acted as biased ago-allosteric modulators, triggered ERK_1/2_ phosphorylation and to a lesser extent, IP_1_ production, stimulated cell proliferation yet were devoid of contractile activity. Interestingly, both hUT-derived pepducins were able to modulate hUII- and URP-mediated contraction albeit to different extents. These new derivatives represent unique tools to reveal the intricacies of hUT signaling and also a novel avenue to design allosteric ligands selectively targeting UT signaling that could prove to be useful for the treatment of hUT-associated diseases.

## Introduction

In humans, the urotensinergic system, composed of a class 1A G protein-coupled receptor (hUT), and two endogenous peptide ligands, urotensin II (UII; hUII = H-Glu-Thr-Pro-Asp-c[Cys-Phe-Trp-Lys-Tyr-Cys]-Val-OH) and urotensin II-related peptide (URP, H-Ala-c[Cys-Phe-Trp-Lys-Tyr-Cys]-Val-OH), continues to represent a promising contemporary target for the treatment of several pathologies (1,2). Notably, multiple studies in animal models have suggested that UT antagonists may represent potential therapeutic agents for treating atherosclerosis (3-6), pulmonary arterial hypertension (7-9), metabolic syndrome (4), and heart failure (10-12). However, in spite of such promise, clinical studies of UT antagonist drug candidates have had limited success due to a lack of efficacy in humans (1,2,13,14). Our current knowledge is therefore insufficient to clearly assess the therapeutic potential of this system and accordingly, a deeper understanding of UT pharmacology is critically needed to accelerate the development of new UT ligands exhibiting efficacy in humans.

While the two endogenous ligands share a common bioactive core, the distinct N-terminal domain of UII isoforms appears to be involved in specific topological changes associated with UT activation (2,15-17). Contingent on their interactions with UT, UII and URP probably induce distinct UT conformational and dynamic changes that lead to divergent signaling profiles with both common and distinct biological activities (2,17-21). Recent years have witnessed the emergence of useful molecules, with probe-dependent actions, that could shed light on the respective roles and importance of UII and URP under normal and pathological conditions (13,17,18,22-27).

Promoting specific GPCR signaling events with biased agonists or allosteric modulators is a potentially innovative way to treat numerous conditions including cardiovascular disease, diabetes as well as neuropsychiatric/neurodegenerative disorders (28). Hence, this notion has been introduced into various drug discovery programs through theoretical predictions based on known signaling components of cells and from studies in knockout animals (reviewed in (29)). However, there are numerous instances where it is still not yet possible to predict what type of signaling bias represents a superior therapeutic approach. In these cases, empirical testing of exemplar molecules in animal models is a way forward. Such tools are currently unavailable in the context of the urotensinergic system.

Pepducins are lipidated cell-penetrating peptides composed of a lipid moiety attached to a peptide corresponding to an amino acid segment from one of the cytoplasmic loops of a G protein-coupled receptor (GPCR) of interest (reviewed in (30,31)). Following establishment of an equilibrium between the inner and outer leaflets of the lipid bilayer, pepducins interact with their cognate GPCR leading to stabilization of a restricted subset of its inactive and active conformational states (32-34). Hence, such compounds can function as allosteric agonists or positive/negative allosteric modulators, making them useful for the study of GPCR signaling, as reported for protease-activated receptors (33,34), chemokine receptors (35,36), and β-adrenergic receptors (37), as well as for the potential treatment of various diseases including inflammatory diseases, cardiovascular pathologies, and cancer (38).

Herein, we describe the design and pharmacological characterization of two pepducins derived from the second (hUT-Pep2) and third ([Trp^1^, Leu^2^]hUT-Pep3) intracellular loops of hUT. Our results demonstrate that both hUT-derived pepducins, while non cytotoxic, trigger ERK_1/2_ phosphorylation and IP_1_ accumulation. Using BRET-based biosensors, we observed that UT-Pep3 induced G_q_, G_i_, β-arrestin 1, β-arrestin 2 activation, and epidermal growth factor receptor (EGFR) transactivation while UT-Pep2 only activated G_i_, β-arrestin-2 and EGFR transactivation. Interestingly, while only [Trp^1^, Leu^2^]hUT-Pep3 was able to induce proliferation in HEK 293-hUT cells, both pepducins were able to trigger proliferation of neonatal rat cardiac fibroblasts. Further, while both hUT-derived pepducins are unable to trigger rat aortic ring contraction on their own, they could modulate hUII- and URP-mediated contraction to different extents. These new molecular tools represent unique UT-targeted ligands that can be used to interrogate the involvement of specific pathways in hUT-associated diseases or to develop pharmacological agents with fewer side effects and a unique and more precise action for the treatment of various pathologies.

## Results

### Design and toxicity testing of hUT-derived pepducins

Pepducins are composed of a synthetic peptide, mimicking an intracellular GPCR loop, to which a hydrophobic moiety, most commonly the fully saturated C_16_ fatty acid palmitate, is conjugated at their N-terminus (39). Based on a predicted structure for hUT (14,15), pepducins corresponding to the sequence of hUT intracellular loops 2 and 3 (Table 1) were synthesized using solid phase peptide synthesis. hUT-Pep2 is identical to the ICL2 sequence found in hUT. The second pepducin ([Trp^1^, Leu^2^]hUT-Pep3) comprises the hUT ICL3 sequence; however, the first two N-terminal amino acids have been replaced by the corresponding residues from rat UT ICL3 (-Arg-Arg-in hUT and -Trp-Leu-in rUT). This modification was necessary since the pepducin derived from hUT ICL3, comprising at least 7 positively charged residues, was highly cytotoxic, which prevented further use. Nonetheless, as shown in Figure 1A and 1B, hUT-Pep2 and [Trp^1^, Leu^2^]hUT-Pep3 (10^-5^ M) did not significantly affect HEK 293 cell viability or promote LDH release into culture media, thus demonstrating that both pepducins did not reduce cell viability.

**Figure 1.**
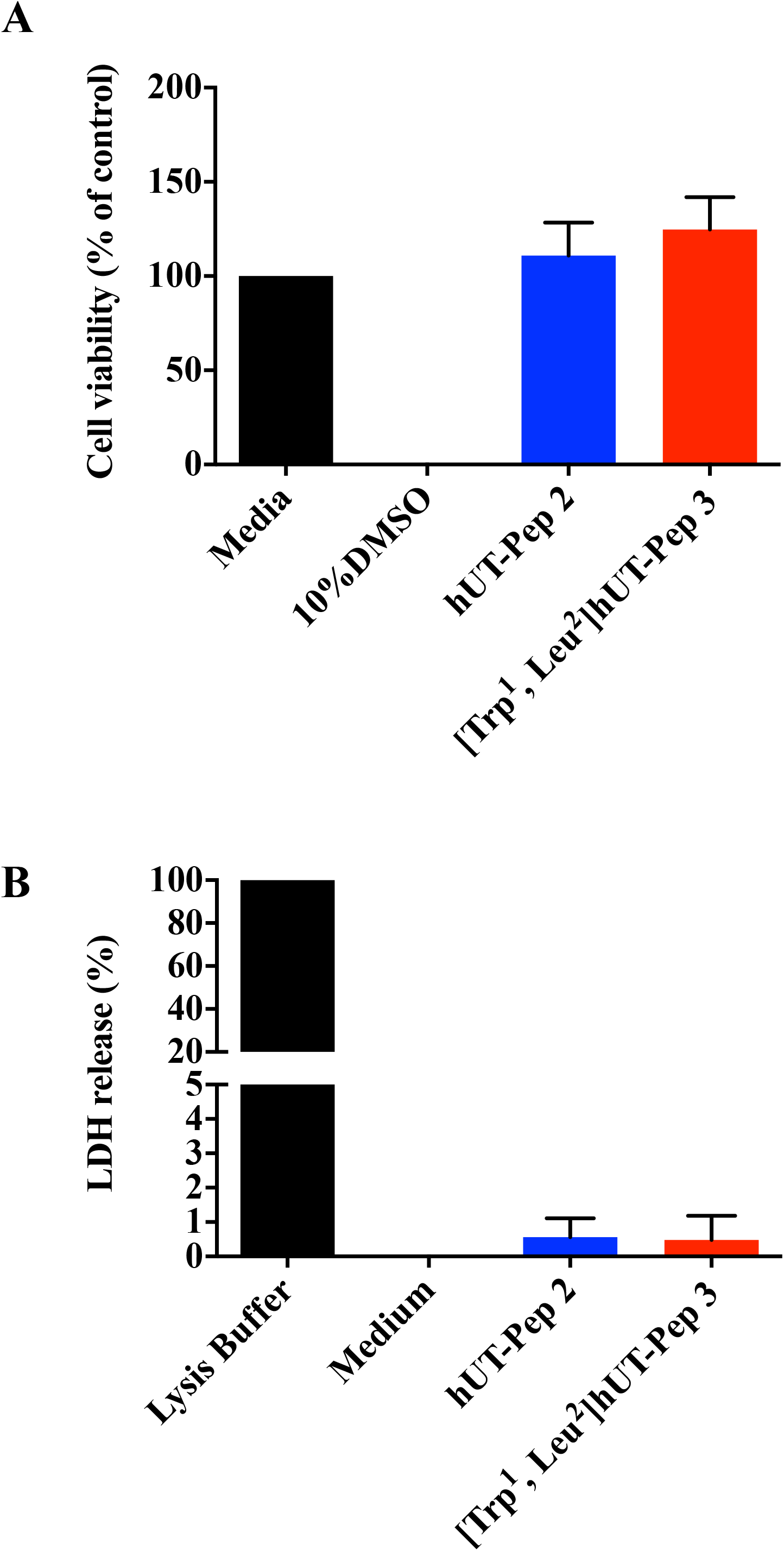
Cytotoxic action of UT-pepducins. Toxicity of UT-Pep2 and UT-Pep3 were evaluated using (A) MTT or (B) LDH assays in HEK 293-*h*UT cells

**Table 1.** Amino acid sequences and analytical data of compounds hUII, URP, and hUT-derived pepducins.

| Cpd Name | Sequence | MW<sup>a</sup> calc | MS<sup>b</sup> found |
| --- | --- | --- | --- |
| hUII | H-Glu-Thr-Pro-Asp-[Cys-Phe-Trp-Lys-Tyr-Cys]-Val-OH | 1387.6 | 1388.1 |
| URP | H-Ala-[Cys-Phe-Trp-Lys-Tyr-Cys]-Val-OH | 1016.4 | 1016.4 |
| hUT-Pep2 | Palm-Arg-Pro-Leu-Asp-Thr-Val-Gln-Arg-Pro-Lys-Gly-Tyr-NH<sub>2</sub> | 1666.0 | 1667.1 |
| [Trp<sup>1</sup>, Leu<sup>2</sup>]hUT-Pep3 | Palm-Trp-Leu-Ser-Gln-Arg-Ala-Ser-Phe-Lys-Arg-Ala-Arg-Arg-NH<sub>2</sub> | 1899.3 | 1899.9 |

<sup>a</sup> Molecular weight calculated on a CEM Liberty Blue peptide synthesizer software. <sup>b</sup> MALDI-TOF mass spectral analysis (m/z). Values represent the observed m/z of the monoisotope [M + H]<sup>+</sup> ions. Palm: palmitoyl.

### hUT-Pep2 and [Trp^1^, Leu^2^]hUT-Pep3 conserve a structure similar to the one adopted within the fulllength receptor

Conformational analysis of hUT-Pep2 and [Trp^1^, Leu^2^]hUT-Pep3 was performed by solution NMR in DPC micelle suspensions, a solvent system commonly used in NMR and CD studies to mimic the zwitterionic membrane environment (40). hUT-Pep2 gave well-resolved NMR spectra and many NMR parameters indicated a folded structure (Tables S1 and S2). In particular, medium-range Nuclear Overhauser effects (NOEs, Table S2) between Hα(i) and HN(i+2) pointed to β-turn structures confirmed by other diagnostic parameters, such as amide temperature coefficients and ^3^*J*_HN-HA_ coupling constants. Actually, 135 NOEs (Table S2) were used in structure calculation, which corresponds to 11.2 NOEs per residue, a ratio similar to what is found in structured protein (41). Among those, 22 medium range NOEs in line with folded (β-turn) peptides were observed. Figures of the NOESY spectra of hUT-Pep2 and [Trp^1^, Leu^2^]hUT-Pep3 are shown in the supporting information (Figure S1 and S2). Restrained MD calculations provided the well-defined structures shown in Figure 2A. Starting from the N-terminus, two type I β-turns along residues 1-4 and 3-6, and three distorted (type IV) β-turns along residues 4-7, 5-8 and 6-9 can be observed in the simulated structures of hUT-Pep2. Interestingly, the obtained structure can be easily overlapped with the corresponding ICL2 segment of a hUT model recently obtained by us (Figure 2B) (15). hUT-Pep2 therefore preserves the conformational propensities of the corresponding ICL2 segment embedded in the wild type receptor. Unfortunately, NMR spectra of [Trp^1^, Leu^2^]hUT-Pep3, despite complete assignment of all the proton resonances (Table S3), showed many broad and overlapping signals particularly in the amide proton region that prevented the measure of many NMR parameters (^3^*J*_HN-HA_ coupling constants and most of the NOEs), and therefore NMR-based calculation of its 3D structure. However, retrievable NMR parameters from the spectra (a few NOEs, up-field shifts of the Hα resonances, and amide temperature coefficients, Tables S3 and S4) suggest a high tendency of [Trp^1^, Leu^2^]hUT-Pep3 to fold into a helix. Using circular dichroism (CD) analyses, we were able to observe that [Trp^1^, Leu^2^]hUT-Pep3, under any condition tested, was able to assume structural motifs (Figure 2C). In 0.1 mg.mL^-1^ SDS solution (red) it appeared to assume a β-sheet conformation (lower peak at 216 nm), while in TFE solutions it appeared to fold as an a-helix (dark green, TFE 80%, lower peaks at 205 and 216 nm). In 0.5 mg.mL^-1^ DPC solution, it appeared to assume a less structured a-helix conformation, which correlates with our NMR data, while in water a non-structured shape (random coil) was adopted. Worth noting, a pepducin derived from the ICL3 of PAR1 studied in DPC solutions, also showed a stable helical conformation (42). Together, these results suggest that both pepducins probably adopt conformations closely related to those found within the wild type receptor.

**Figure 2.**
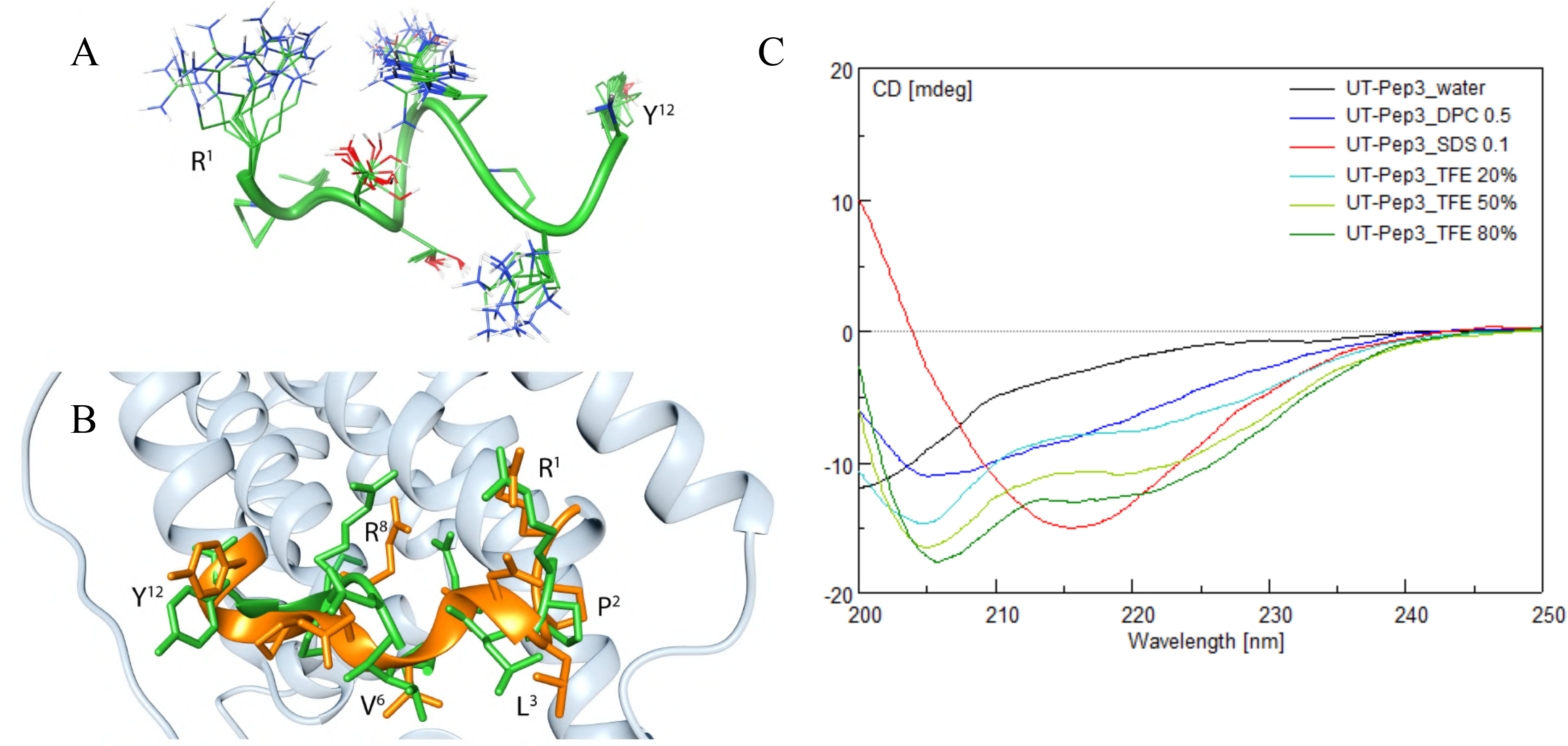
3D structural data of UT-derived pepducins. **(A)** Modeled structure of the hUT-Pep2. Ensemble of the 10 lowest energy conformers of hUT-Pep2. Structures were superimposed using the backbone heavy atoms. Side chain heavy atoms are shown with different colours (carbon, green; nitrogen, blue; oxygen, red). Many hydrogen atoms and the lipid chains are not shown for clarity. Backbone structure is represented as green ribbon. **(B)** Superposition of hUT-Pep2 structure (representing the NMR lowest energy conformer, green) and the second intracellular loop of a hUT model (orange). **(C)** Circular dichroism of [Trp^1^, Leu^2^]hUT-Pep3 in various solvents.

### hUT-derivedpepducins trigger IP_1_ production

To assess hUT-mediated activation of phospholipase C (PLC), the inositol 1,4,5-trisphosphate metabolite inositol monophosphate (IP_1_) was quantified. In agreement with previously reported data (43), we observed that hUII peptide induced a time-depenent increase in IP_1_ that reached a plateau after 40 min (Figure 3A). Concentration-response curves, constructed after 40 min incubation, revealed that hUT-Pep2 and [Trp^1^, Leu^2^]hUT-Pep3 induce IP_1_ production in CHO cells stably expressing hUT (Figure 3B) but not CHO mock-transfected cells (data not shown). Compared to hUII, hUT-Pep2 and [Trp^1^, Leu^2^]hUT-Pep3 appeared to be less potent (pEC_50_ < 6) at promoting IP_1_ production (Figure 3B), acting as very weak allosteric agonist.

**Figure 3.**
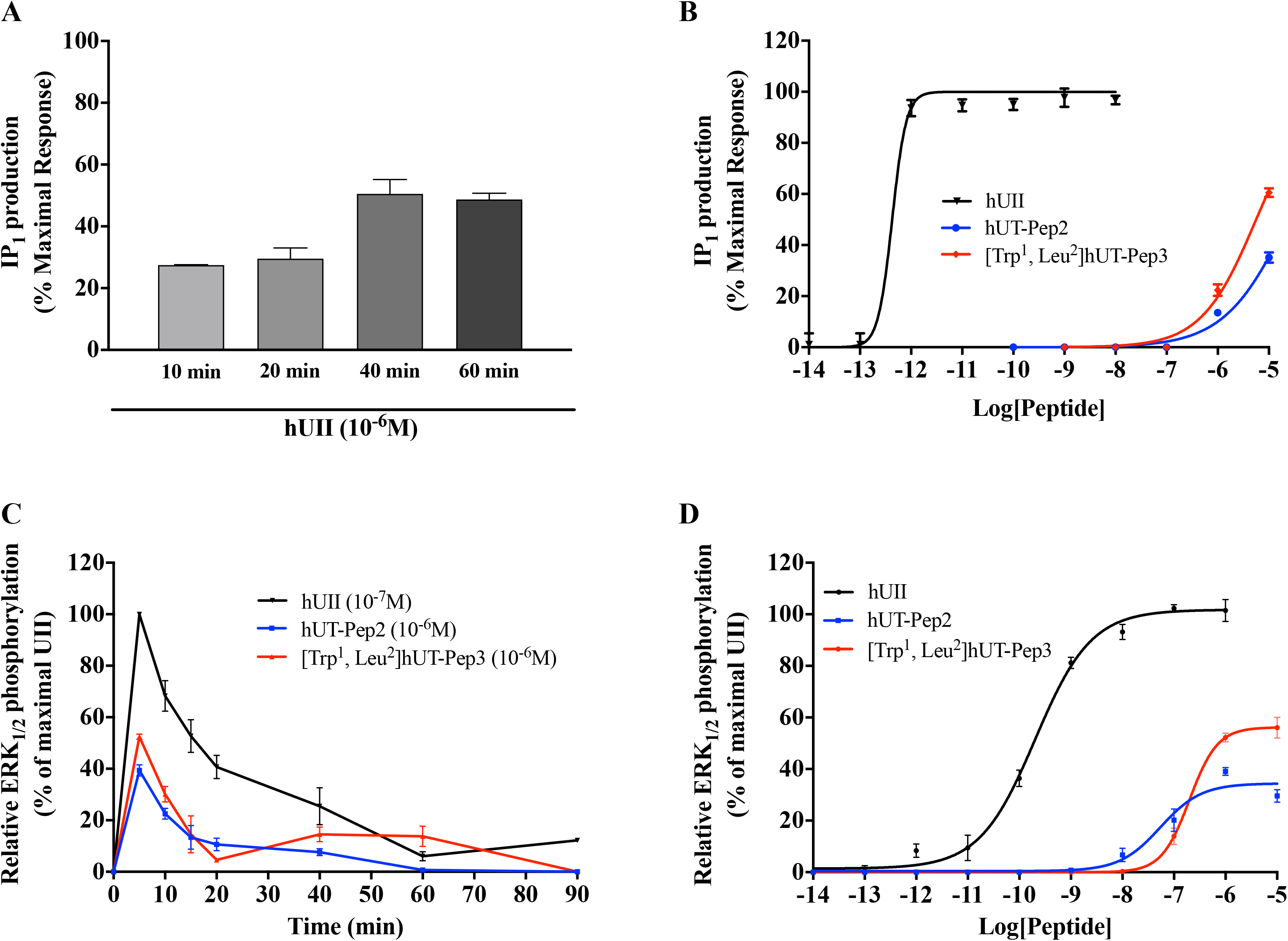
IP_1_ (A and B) and ERK_1/2_ phosphorylation (C and D) kinetics and concentration-dependent stimulation following hUT receptor activation. IP_1_ production was evaluated using the IP-One terbium immunoassay kits while ERK_1/2_ phosphorylation was measured by western blot in CHO-K1 cells stably expressing the hUT receptor. The data are normalized to the hUII maximum effect. Each curve represents the mean ± SEM of at least three independent experiments performed in triplicate.

### hUT-derived pepducins trigger ERK_1/2_ phosphorylation through different mechanisms

Kinetic measures, using the CHO-hUT cell line (17), revealed that both hUT-derived pepducins induced ERK_1/2_ phosphorylation. Used at 10^-6^ M, hUT-Pep2 and [Trp^1^, Leu^2^]hUT-Pep3 exerted their maximal effect at 5 min, similar to hUII (Figure 3C). While a second peak was noticeable for UT-Pep3 at 60 min, a similar pattern was not observed for hUT-Pep2, suggesting a potential signaling bias (Figure 3C). We next generated concentration-response curves for each pepducin and hUII (Figure 3D). Compared to hUII, hUT-Pep2 and [Trp^1^, Leu^2^]hUT-Pep3 were both less potent (pEC_50_ = 7.26 ± 0.14 and 6.70 ± 0.06, respectively) and less efficacious (E_max_ = 34 ± 2% and 56 ± 2%, respectively) at triggering ERK_1/2_ phosphorylation (Figure 3D), behaving as weak partial allosteric agonists.

As shown in Figure 4A, in the absence of the receptor, neither hUII nor hUT-derived pepducins were able to trigger ERK_1/2_ phosphorylation, therefore demonstrating their specificity for hUT. Following pre-treatment of CHO-hUT cells with PTX (a G_i/o_ protein inhibitor), U73122 (a broad range PLC inhibitor), or AG1478 (a specific EGFR inhibitor), ERK_1/2_ phosphorylation was evaluated at 5 min as described above. While PTX and U73122 completely prevented [Trp^1^, Leu^2^]hUT-Pep3-mediated ERK_1/2_ phosphorylation, hUT-Pep2 action was only partially affected by the PLC inhibitor, further supporting distinct mechanisms of action (Figure 4B and 4C). Inhibition of EGFR transactivation with AG1478 significantly reduced the ability of both hUT-derived pepducins to trigger ERK_1/2_ phosphorylation (Figure 4D).

**Figure 4.**
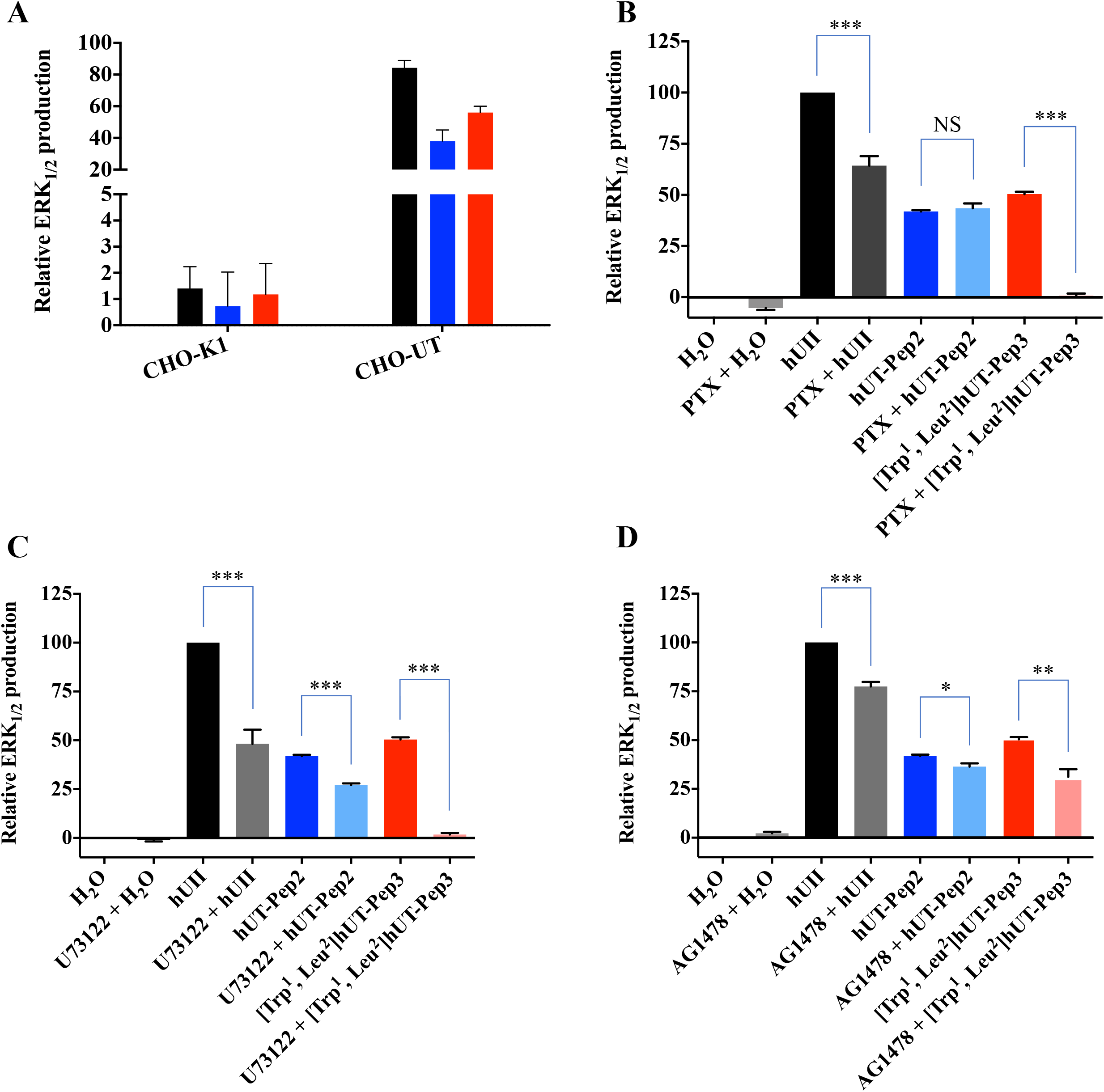
ERK_1/2_ phosphorylation profile after hUT receptor activation. ERK_1/2_ phosphorylation was measured by western blot in CHO-K1 cells or CHO-K1 cells stably expressing the hUT receptor. (A) ERK_1/2_ phosphorylation normalized to total ERK was measured following hUII (10^-7^M), hUT-Pep2, or [Trp^1^, Leu^2^]hUT-Pep3 (10^-6^M) treatments in CHO-K1 or CHO-K1-hUT cells. (B-D) The hUT receptor was activated by hUT-Pep2, or [Trp^1^, Leu^2^]hUT-Pep3 (10^-6^M) under control conditions, or after treatment with PTX, U73122 or AG1478. The data are normalized to the UII maximum effect in B, C and D. Each curve represents the mean ± SEM of at least three independent experiments performed in triplicate.

Consistent with our previous results, kinetic experiments, performed this time using a BRET-based biosensor and HEK 293 cells stably expressing hUT, once again revealed biphasic ERK_1/2_ activation following hUII treatment with a maximum activation after 5 min (Figure 5A). hUT-Pep2 (10^-5^ M) also reached its maximal effect after 5 min while [Trp^1^, Leu^2^]hUT-Pep3 needed 10 min to reach an efficacy quite similar to what was observed with hUII (10^-7^ M). Both hUT-derived pepducins appeared to return to basal levels of ERK_1/2_ activation but with different kinetics: hUT-Pep2 reaching basal levels after 10 min while [Trp^1^, Leu^2^]hUT-Pep3 was still decreasing after 20 min (around 50%) (Figure 5B and 5C). Pre-treatment with YM254890, a selective inhibitor of G_q_ signaling, almost completely abolished the ability of hUII to increase ERK_1/2_ phosphorylation while having a modest, but significant, inhibitory effect on the actions of hUT-Pep2 (at 5 min, \**P* < 0.05) and [Trp^1^, Leu^2^]hUT-Pep3 (at 5 and 10 min, *\*P* < 0.05) (Figure 5A-C). Interestingly, while hUT-Pep2-mediated ERK_1/2_ phosphorylation returned to basal levels after 10 min, we observed that in the presence of YM254890, a second peak is clearly detectable at 20 min (Figure 5B). Similarly, while a small but significant reduction of ERK_1/2_ phosphorylation is observed with [Trp^1^, Leu^2^]hUT-Pep3 in the presence of YM254890, this activation is maintained and did not decrease over the studied period of time (Figure 5C). Finally, and similar to what was observed in CHO-hUT cells using western blotting, hUT-Pep2 and [Trp^1^, Leu^2^]hUT-Pep3 triggered ERK_1/2_ phosphorylation with a lower potency and efficacy than hUII (Figure 6A). Together, these results support a different mode of action related to the regulation of ERK_1/2_ activity by hUT-Pep2 and [Trp^1^, Leu^2^]hUT-Pep3.

**Figure 5.**
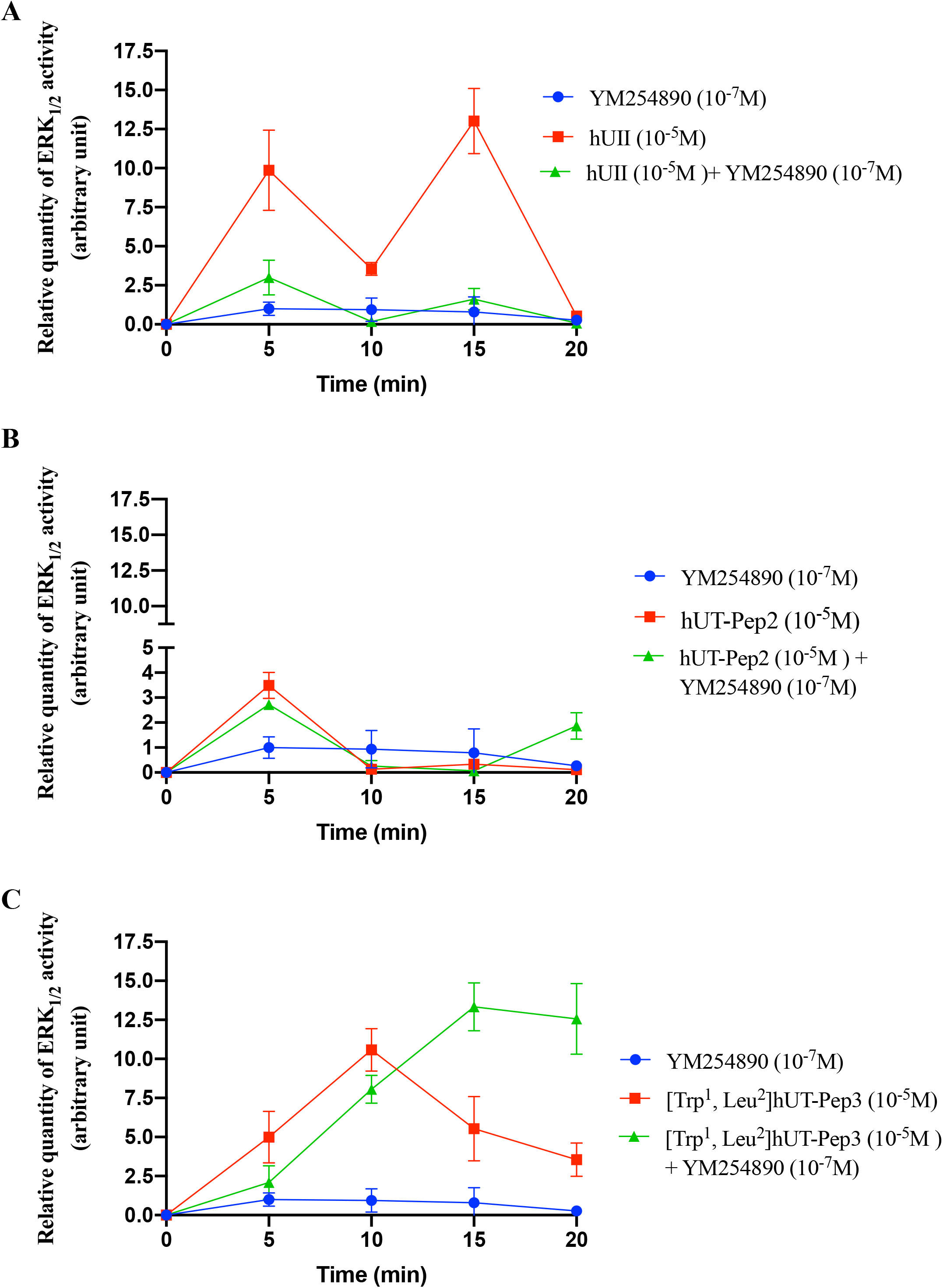
ERK_1/2_ phosphorylation kinetics after hUT receptor activation. Activation of hUT by hUII, hUT-Pep2 or [Trp^1^, Leu^2^]hUT-Pep3 under control conditions or after treatment with YM254890. Data are normalized to the hUII maximum effect. Curves represent means ± sem of 3 independent kinetic experiments performed in triplicate.

**Figure 6.**
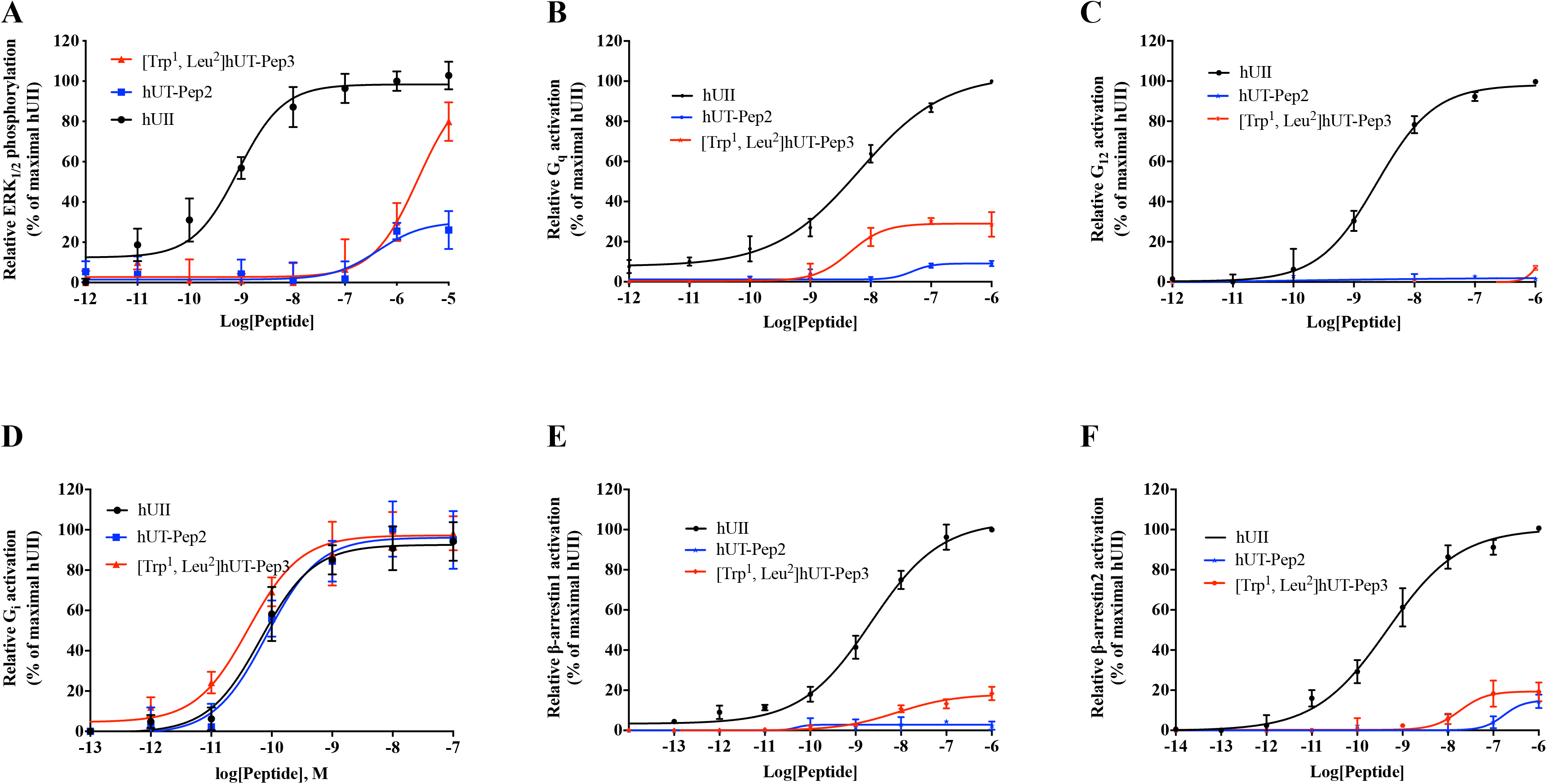
Signaling signature of UT-pepducins following UT receptor activation. BRET measurements were performed following hUT stimulation with increasing concentrations of hUII, or UT-derived pepducins in HEK 293-hUT cells. The data are normalized to maximal hUII responses (10^-6^M). Each curve represents the mean ± SEM of at least three independent concentration-response experiments performed in triplicate.

### Signaling signatures of hUT-derived pepducins

We next investigated the propensity of each pepducin to induce G_q_, G_12_, G_i_ activation or promote conformational changes at β-arrestin 1 and β-arrestin 2 using BRET-based biosensors. As depicted in Figure 6B-D and Table 2, hUT-derived pepducins can promote activation of G_q_, albeit with low efficacy, and G_i_ but were almost completely unable to activate G_12_. Notably, on one hand, [Trp^1^, Leu^2^]hUT-Pep3 stimulated G_q_ activation with a similar potency (pEC_50_ = 8.34 ± 0.33) but a lower efficacy (E_max_ = 29 ± 4%, \*\*\**P* < 0.001) than hUII (pEC_50_ = 8.19 ± 0.12; E_max_ = 109 ± 5%) acting as a partial allosteric agonist of this pathway (Figure 6B; Table 2). On the other hand, hUT-Pep2 behaved as a weak partial allosteric agonist (pEC_50_ = 7.31 ± 0.55) of this pathway reaching approximately 9% of the hUII maximum response (Figure 6B; Table 2). Interestingly, in contrast to hUII, both hUT-derived pepducins acted as full agonists for the activation of G_i_ (Figure 6D; Table 2).

**Table 2.** Signaling signatures of hUII and hUT-derived pepducins

| Peptides | G<sub>q</sub> |  | G<sub>12</sub> |  | G<sub>i</sub> |  | β-arr1 |  | β-arr2 |  |
| --- | --- | --- | --- | --- | --- | --- | --- | --- | --- | --- |
|  | pEC<sub>50</sub><sup>a</sup> | E<sub>max</sub><sup>b</sup> | pEC<sub>50</sub><sup>a</sup> | E<sub>max</sub><sup>b</sup> | pEC<sub>50</sub><sup>a</sup> | E<sub>max</sub><sup>b</sup> | pEC<sub>50</sub><sup>a</sup> | E<sub>max</sub><sup>b</sup> | pEC<sub>50</sub><sup>a</sup> | E<sub>max</sub><sup>b</sup> |
| hUII | 8.19 ± 0.12 | 109 ± 5 | 8.62 ± 0.09 | 98 ± 3 | 10.18 ± 0.07 | 93 ± 2 | 8.66 ± 0.13 | 104 ± 4 | 9.33 ± 0.14 | 100 ± 3 |
| hUT-Pep2 | 7.31 ± 0.55** | 9 ± 1*** | - | - | 10.09 ± 0.08 | 96 ± 2 | - | - | - | 15<sup>c</sup> |
| [Trp<sup>1</sup>, Leu<sup>2</sup>]hUT-Pep3 | 8.34 ± 0.33 | 29 ± 4*** | - | 5<sup>c</sup> | 10.39 ± 0.09 | 97 ± 2 | 8.08 ± 0.27** | 18 ± 2*** | 7.74 ± 0.32*** | 19 ± 3*** |

<sup>a</sup>pEC<sub>50</sub> = -log EC<sub>50</sub>. <sup>b</sup>The maximum efficacy is expressed as a percentage of the activation induced by hUII (10<sup>-6</sup>M). <sup>c</sup>Maximum efficacy at 10<sup>-6</sup>M. All values are expressed as mean ± SEM. Statistical analysis were performed using unpaired Student's t test, \*\**P* ≤ 0.01, \*\*\**P* ≤ 0.001, versus values obtained for hUII. Each replicate (n = 3-5) was conducted on different cellular passages.

We next evaluated the propensity of hUT-derived pepducins to trigger conformational changes in β-arrestin 1 and 2. For β-arrestin 1, [Trp^1^, Leu^2^]hUT-Pep3, which is almost equipotent (pEC_50_ = 8.08 ± 0.27; \*\**P* < 0.01) compared to hUII (pEC_50_ = 8.66 ± 0.13) but only reached around 18% of its maximum response (Figure 6E; Table 2), acting as a weak partial allosteric agonist for this signaling pathway while hUT-Pep2 appears inactive. For β-arrestin 2, [Trp^1^, Leu^2^]hUT-Pep3, once again acted as a weak partial allosteric agonist, triggering activation of this pathway with low potency (pEC_50_ = 7.74 ± 0.32; \*\*\**P* < 0.001) and efficacy (E_max_ = 19 ± 3; \*\*\**P* < 0.001) compared to hUII (pEC_50_ = 9.33 ± 0.14; E_max_ = 100 ± 3%) (Figure 6F; Table 2). hUT-Pep2 behaved as a very weak partial allosteric agonist reaching, at 10^-6^M, around 15% of the maximal response evoked by hUII or URP. Together, these results demonstrate that both pepducins can trigger hUT activation albeit to different extents.

### hUT-derived pepducins promote EGFR transactivation through G_q_ activation

We next investigated whether UT stimulation by hUT-Pep2 or [Trp^1^, Leu^2^]hUT-Pep3 could mediate EGFR transactivation. Untransfected cells and cells stably expressing wild type hUT were transfected with EGFR-GFP and stimulated with either hUII, hUT-Pep2 or [Trp^1^, Leu^2^]hUT-Pep3 (10 ^-6^ M). In the absence of stimulation, EGFR had a uniform membrane distribution in HEK 293 cells stably transfected or not with hUT (Figure 7A and 7B). In contrast, treatment with hUII resulted in EGFR internalization as characterized by a marked redistribution into cellular aggregates compared to untreated HEK 293 cells (Figure 7B and 7E). Treatment with hUT-derived pepducins in HEK 293-hUT cells resulted in significant internalization of EGFR-GFP as demonstrated by the punctate distribution of fluorescence (Figure 7H and 7K), which appeared stronger upon [Trp^1^, Leu^2^]hUT-Pep3 stimulation. Finally, we observed that pretreatment of the cells with the specific G_q_ inhibitor YM254890 almost completely abolished hUII-as well as UT-derived pepducin-associated EGFR transactivation (Figure 7F, 7I, and 7L). No effects of any of the ligands were seen in mock-tranfected cells (Figure 7D, 7G, and 7J). Together, these results revealed that both pepducins can promote EGFR transactivation, albeit to different extents, and that this activation is abrogated following G_q_ inhibition.

**Figure 7.**
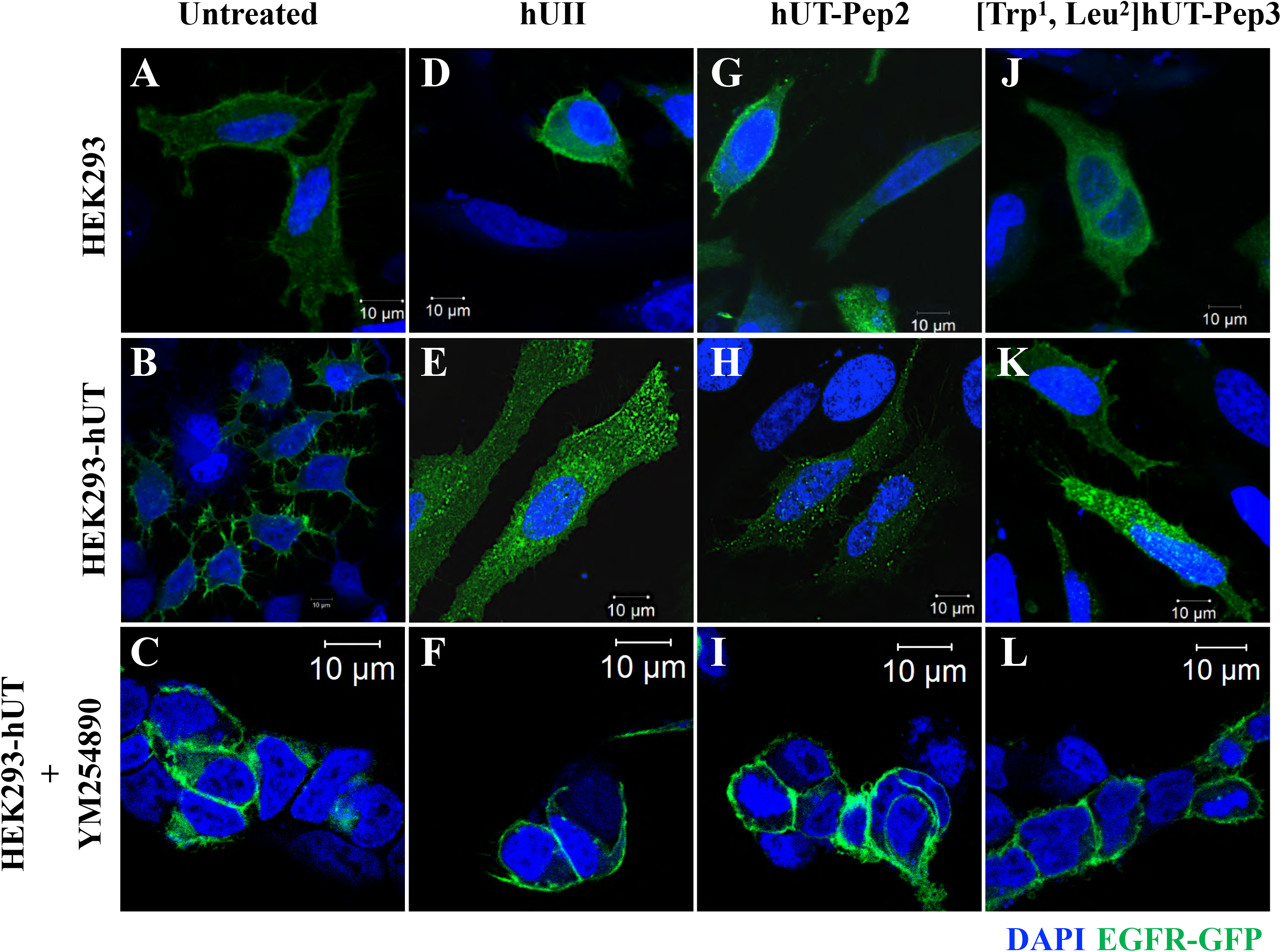
UT-pepducin-mediated transactivation of EGFR following hUT activation. HEK 293 cells stably expressing wild type hUT (B, E, H, K) or not (A, D, G, J) and transfected with FLAG-EGFR were treated with hUII, hUT-Pep2, or [Trp^1^, Leu^2^]hUT-Pep3 in the presence (C, F, I, L) or not of YM254890. EGFR internalization, following hUT stimulation with various ligands for 30 min, was visualized using confocal microscopy. The images shown are from representative experiment repeated at least three times.

### hUT-Pep2 and [Trp^1^, Leu^2^]hUT-Pep3 can induce rat neonatal fibroblast proliferation

Several studies have demonstrated the mitogenic properties of UII, URP and related analogs in various cultured cells (1), including rat fibroblasts (44,45), and astrocytes (20), as well as HEK 293-hUT cells (43). In the present study, using HEK 293-hUT cells and two different methods, *i.e.* MTT assays and flow cytometry, we observed that hUII and [Trp^1^, Leu^2^]hUT-Pep3 but not hUT-Pep2 were able to induce HEK 293-hUT cell proliferation (Figure 8A and 8C). Again, in the absence of the receptor, the above described effect was completely abrogated (Figure 8B). Interestingly, using high content imaging, we observed that both hUT-derived pepducins produced a proliferative action in rat neonatal fibroblasts similar to what was observed with hUII (Figure 8D). These observations showed that hUT-Pep2 and [Trp^1^, Leu^2^]hUT-Pep3 can act as allosteric agonists promoting cell proliferation and that this action is cell-context dependent, at least for hUT-Pep2.

**Figure 8.**
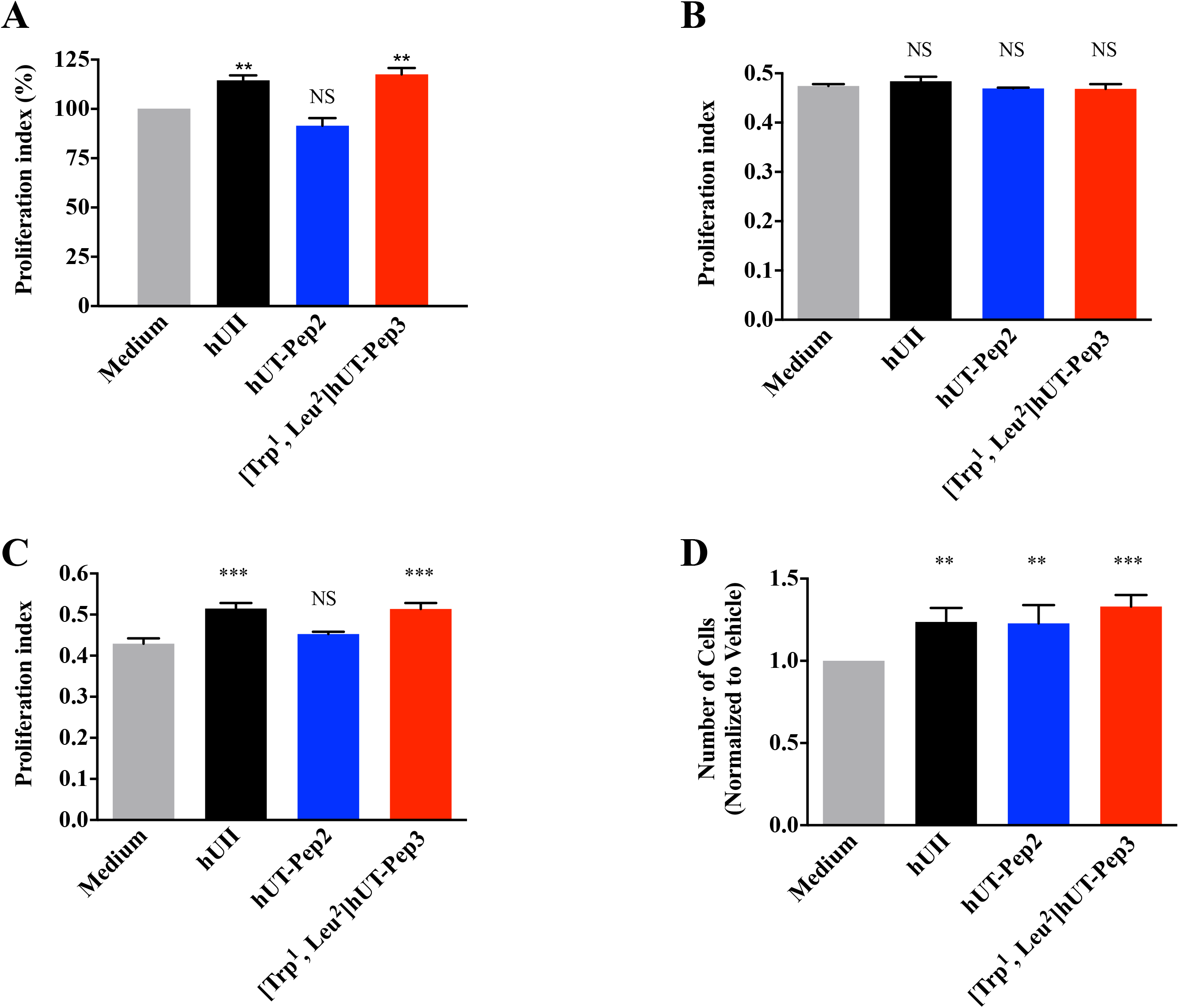
Effects of hUII and UT-derived pepducins on proliferation of hUT receptor-expressing HEK 293 cell and rat neonatal fibroblasts. HEK 293 cells stably expressing hUT were treated with hUII, hUT-Pep2 or [Trp^1^, Leu^2^]hUT-Pep3 (10^-6^M) and cell proliferation was evaluated by MTT assay **(A)** or flow cytometry **(C)**. HEK 293 cells were treated with hUII, hUT-Pep2 or [Trp^1^, Leu^2^]hUT-Pep3 (10^-6^M) and cell proliferation was also evaluated by MTT assay **(B)**. **(D)** Rat neonatal fibroblasts endogenously expressing UT were treated with hUII, hUT-Pep2 or [Trp^1^, Leu^2^]hUT-Pep3 (10^-6^M) and proliferation was evaluated with an Opera Phenix using smooth muscle a-actin as primary antibody and an anti-rabbit Alexa Fluor 488 as secondary antibody. Each bar corresponds to the mean ± SEM obtained from 3 independent experiments conducted in triplicate.

### hUT-derivedpepducins can block hUII- and URP-mediated contraction

Since its discovery, UII and its paralog peptide URP have been considered potent vasoconstrictors (1). As shown in Figure 9A, none of the hUT-derived pepducins were able to stimulate contraction reaching only between 5 to 15% of the force induced by hUII or URP in rat aortic ring preparations (Table 3). Used as potential allosteric modulators, hUT-Pep2 and [Trp^1^, Leu^2^]hUT-Pep3 suppressed, albeit to different extents, the maximum contractile response to hUII and URP (Figure 9B and 9C; Table 4). For instance, pre-treatment with [Trp^1^, Leu^2^]hUT-Pep3 at 10^-6^ M, produced significant suppression of the maximum contractile response to hUII (E_max_ = 35 ± 6%; \*\*\**P* < 0.001) and URP (E_max_ = 16 ± 2%; \*\*\**P* < 0.001) (Table 3). Similarly, hUT-Pep2 at the same concentration, promoted a significant but small reduction of the maximum contractile response to hUII (E_max_ = 86 ± 3%; \**P* < 0.05) and URP (E_max_ = 76 ± 8%; \*\**P* < 0.01). Interestingly, hUT-Pep2 induced a significant rightward shift in the concentration-response curve of hUII but not URP, therefore suggesting a probe-dependent action. These data demonstrate that hUT-Pep2 but most importantly [Trp^1^, Leu^2^]hUT-Pep3 can negatively modulate hUII- and URP-mediated contraction.

**Figure 9.**
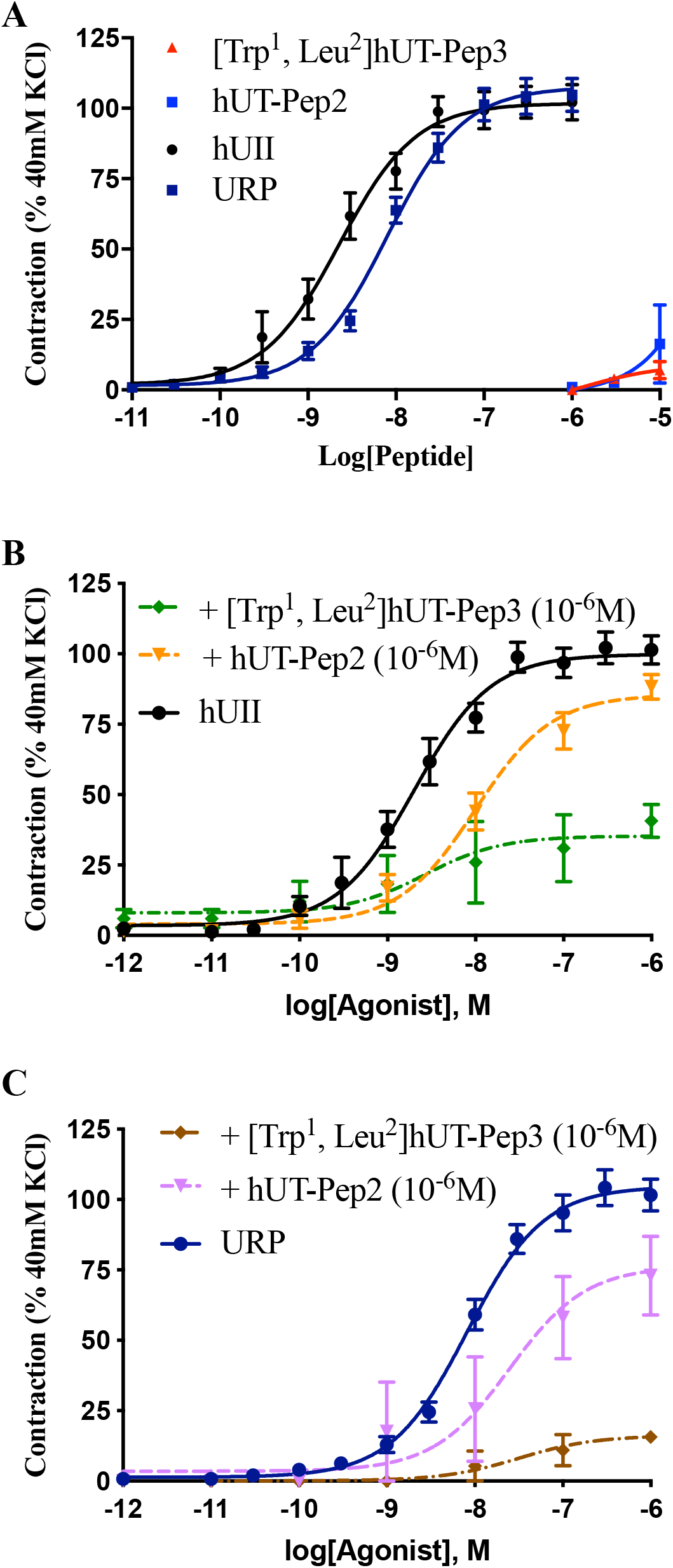
Vasocontractile action of UT-derived pepducins. (A) Representative concentration-response curves obtained with rat thoracic aortic rings after adding cumulative concentrations of hUII or UT-derived pepducins. Representative concentration-response curves obtained with rat thoracic aortic rings after adding cumulative concentrations of hUII (B) or URP (C) following pre-treatment with hUT-Pep2 or [Trp^1^, Leu^2^]hUT-Pep3. Each replicate (n) was conducted on tissue from at least 3 different animals. Data represent mean ± S.E.M.

**Table 2.** Contractile activity of UT-derived pepducins.

| | $E_{\max}^a$ | $pEC_{50}^b$ | n |
| --- | --- | --- | --- |
| hUII | $102 \pm 3$ | $8.64 \pm 0.09$ | 22 |
| URP | $108 \pm 3$ | $8.10 \pm 0.06$ | 22 |
| hUT-Pep2 | 7% @ 10μM | <5 | 5 |
| [Trp<sup>1</sup>, Leu<sup>2</sup>]hUT-Pep3 | 16% @ 10μM | <5 | 5 |

<sup>a</sup>Maximum efficacy is expressed as a percentage of the amplitude of the contraction induced by KCl (40 mM). All values are expressed as mean ± SEM. Each replicate (n) was conducted on tissue from different animals.

**Table 4.** Antagonist contractile activities.

| *Peptide* | Aortic ring contraction hUII |  |  | Aortic ring contraction URP |  |  |
| --- | --- | --- | --- | --- | --- | --- |
|  | n | pEC<sub>50</sub> | E<sub>max</sub><sup>a</sup> | n | pEC<sub>50</sub> | E<sub>max</sub><sup>a</sup> |
| hUII | 19 | 8.69 ± 0.08 | 100 ± 3 |  |  |  |
| URP |  |  |  | 22 | 8.09 ± 0.06 | 105 ± 3 |
| hUT-Pep2 | 5 | 8.00 ± 0.11** | 86 ± 3* | 5 | 7.60 ± 0.31 | 76 ± 8** |
| [Trp<sup>1</sup>, Leu<sup>2</sup>]hUT-Pep3 | 5 | 8.57 ± 0.63 | 35 ± 6*** | 5 | 8.59 ± 0.12 | 16 ± 2*** |

Aortic rings were pre-treated with different analogues at a concentration of 10<sup>-6</sup>M for 30 minutes prior to hUII or URP treatment. <sup>a</sup>Maximum efficacy is expressed as a percentage of the KCl-induced contraction (40 mM) divided by the tissue-response induced by hUII or URP (10<sup>-6</sup>M). All values are expressed as mean ± SEM. Statistical analysis were performed using unpaired Student's t test, \*P ≤ 0.05, \*\*P ≤ 0.01, \*\*\*P ≤ 0.001, versus values obtained for hUII or URP. Each replicate (n) was conducted on tissue from different animals.

## Discussion

By selectively engaging only a subset of potential intracellular partners, allosteric agonists or biased ligands may offer an unparalleled means to understand and control GPCR-mediated signaling (46). Over the years, the pepducin concept has yielded numerous allosteric agonists and antagonists for several GPCRs, some of them reaching clinical trials (47-49). By stabilizing or disrupting molecular interactions that change the energy landscape of a GPCR, pepducins have the potential to affect the conformational ensemble in ways that affect signaling (50). In the present study, we tested the ability of pepducins, derived from the sequence of intracellular loops 2 and 3 of hUT, to act as functionally-selective modulators. Using various BRET-based biosensors, we demonstrated that hUT-Pep2 and [Trp^1^, Leu^2^]hUT-Pep3 showed a clear bias toward the activation of G_i_, compared to G_q_ or G_12_ activation as well as β-arrestin conformational changes. Most notably, these lipidated peptides, which assumed a 3-dimensional structure similar to those adopted in the full-length receptor and did not show any cytotoxic effects, were both able to act as allosteric biased agonists by promoting hUT-related signaling. Compared to hUII, hUT-Pep2 and [Trp^1^, Leu^2^]hUT-Pep3 were able to stimulate, albeit with lower potency and efficacy, ERK_1/2_ phoshorylation through different mechanisms. hUT-Pep2-induced ERK_1/2_ phosphorylation was partially reduced following treatment of the cells with inhibitors of PLC, G_q_ and EGFR, but unaffected by PTX treatment while the actions of hUII were substantially reduced, or even suppressed, under similar conditions. Therefore, G_i_ activation does not appear to mediate the effects of hUT-Pep2 on ERK_1/2_ phosphorylation, which likely instead involves G_q_, β-arrestin2 as well as EGFR transactivation. Conversely, the drastic reduction in [Trp^1^, Leu^2^]hUT-Pep3-induced ERK_1/2_ phosphorylation following inhibition of G_i_ or PLC suggested, at first, a possible involvement of G_i_ and G_q_ in ERK_1/2_ phosphorylation associated with hUT activation by [Trp^1^, Leu^2^]hUT-Pep3. However, G_q_ inhibition resulted in only a small, but significant, reduction in the early phase of ERK_1/2_ phosphorylation with a late phase that was both enhanced and sustained (**Figure 5C**). As previously reported, inhibition of phospholipase C by U73122 can also block receptor-mediated activation of G_i_ proteins (51,52). GPCRs are known to activate PLC, either *via* GTP-liganded α subunits of the G_q_ class of G proteins or by Gβγ dimers liberated from G_i_ proteins (53). Thus, the complete inhibition of [Trp^1^, Leu^2^]hUT-Pep3-mediated ERK_1/2_ phoshorylation by either PTX or U73122 suggests a direct connection between the two that might involve Gβγ subunits of G_i_. In support of this hypothesis, it was demonstrated that dually coupled GPCRs require cooperation of G_i_- and G_q_-mediated pathways for efficient stimulation of the ERK_1/2_ cascade (54). Several studies have suggested that EGFR transactivation plays a critical role in the hypertrophic response evoked by UT in rat cardiomyocytes (55). Interestingly, such activation was prevented by knockdown of β-arrestins (55). However, as demonstrated, the absence of G protein activation prevents β-arrestin signaling (56). To dissect the mechanism of hUII, hUT-Pep2, and [Trp^1^, Leu^2^]hUT-Pep3-induced EGFR translocation, we transfected HEK 293-hUT cells with GFP-EGFR and monitored internalization by confocal microscopy. When stimulated with hUII or a hUT-derived pepducins, GFP-EGFR was redistributed from the plasma membrane into endocytic vesicles. Pre-treatment with YM254890 significantly, if not completely, blocked hUII-, hUT-Pep2- and [Trp^1^, Leu^2^]hUT-Pep3-induced EGFR internalization, indicating a requirement for G_q_ in their respective responses. Based on previous publications (55), it seems that, at least for hUII, EGFR internalization requires both G_q_ and β-arrestins.

Following characterization of their signaling signatures in HEK 293-hUT cells, we next investigated the propensity of hUII-, hUT-Pep2 and [Trp^1^, Leu^2^]hUT-Pep3 to promote cell proliferation in heterologous and native cell lines and trigger/block aortic ring contraction. As previously reported, downstream signaling linked to the mitogenic HEK 293-hUT responses obtained following UII treatment involved ERK_1/2_ activation and the RhoA/Rho kinase pathway (43). In the present work, we noted that hUII and [Trp^1^, Leu^2^]hUT-Pep3, but not hUT-Pep2, were able to trigger HEK 293-hUT cell proliferation. Since both hUT-derived pepducins were unable to trigger G_12_ activation, differences in the capacity of hUT-Pep2 and [Trp^1^, Leu^2^]hUT-Pep3 to exert mitogenic actions could arise, at least in part, from their significantly different efficacy to stimulate ERK_1/2_ phosphorylation, with [Trp^1^, Leu^2^]hUT-Pep3 acting as a weak full agonist while hUT-Pep2 behaved as a partial agonist (**Figure 6A**). As previously demonstrated, hUII can induce development of cardiac remodeling by promoting fibroblast mitogenesis (44). Interestingly, and similar to what was observed with hUII, both hUT-derived pepducins promoted proliferation of rat neonatal cardiac fibroblasts. Such differences in the mitogenic action of hUT-derived pepducins on HEK 293-hUT cells and cardiac fibroblasts might be inferred from their different cellular contexts, or differences between receptor isoforms. Indeed, it is now widely accepted that GPCRs can adopt multiple conformations upon ligand binding and that structurally distinct ligands can stabilize functionally distinct conformations (57). However, parameters that influence the mechanisms and the timescales by which ligands induce/stabilize specific conformations will most likely differ depending on cellular context because of disparities in associated signaling proteins and the composition of the lipid bilayer (58,59). Also, despite a similar tissue distribution, structural differences between rat and human UT isoforms (only 75% sequence similarity while rat and mouse UT share more than 95% homology) as well as distinct patterns of putative phosphorylation and palmitoylation sites probably lead to differences in signaling, internalization and desensitization patterns (2). Such differences could therefore also explain the distinct biological activities observed with hUT-Pep2 in HEK 293-hUT cells and rat neonatal fibroblasts. Finally, both endogenous ligands of the urotensinergic system are known as potent vasoconstrictors (1). In the present study, neither UT-derived pepducin exhibited significant contractile action in rat aortic ring preparations. Since aortic ring contraction is in part mediated by intracellular calcium mobilization following activation of G_q_ (22,23,43) and by the stimulation of the small GTPase RhoA following G_12_ activation (60), the weak contractile potency of hUT-Pep2 and [Trp^1^, Leu^2^]hUT-Pep3 could be explained by their inability to trigger sustained G_12_ and/or G_q_ activation. This is perhaps not surprising as receptor expression is low. We therefore hypothesized that these pepducins, which mostly act as weak agonists in overexpressing cells, could behave as modulators of hUII and URP in an endogenous context. We therefore tested whether agonist-stimulated aortic ring contraction was inhibited by either hUT-Pep2 and/or [Trp^1^, Leu^2^]hUT-Pep3. Interestingly, both pepducins were able to reduce hUII- and URP-mediated contraction albeit to different extents. Their mechanisms of action will need further study. However, and as proposed for other reported pepducins, pepducins acting as negative modulators, could 1) compete with their homologous peptide segments, buffering/hindering the interaction of signaling partners and/or components of signaling cascade or 2) interact with complementary regions of the peptide segment, *i.e.* intracellular loops or transmembrane domains, therefore destabilizing (antagonism) receptor conformations and dynamics ultimately modulating associated signaling (33).

## Conclusions

Taken together, our results demonstrate that hUT-Pep2 and [Trp^1^, Leu^2^]hUT-Pep3, derived from the intracellular loops 2 and 3, respectively, of hUT can act as biased allosteric agonists/antagonists, engaging/preventing different signaling pathways following UT activation. Given that pepducins are believed to act intracellularly by binding directly to targeted GPCRs on the inner face of the membrane (50,61), it is not surprising to observe that different subsets of downstream effectors than those promoted by the binding of hUII are engaged. Interestingly, while hUT-Pep2 and [Trp^1^, Leu^2^]hUT-Pep3 can trigger cell proliferation, this effect appears to be dependent on cellular context. The precise mechanisms of action by which hUT-derived pepducins can promote G_q_/G_i_ activation, EGFR transactivation and fibroblast proliferation are still unknown and will need further investigation. These new molecular tools could represent innovative UT-targeted modulators with unique pharmacological profiles.

## Experimental procedures

### Cell lines and reagents

HEK 293 cells were purchased from ATCC while the stably transfected HEK 293-hUT cell line was generated in our laboratory (23). Cell culture reagents were obtained from Invitrogen while antibodies used to evaluate ERK_1/2_ phosphorylation, *i.e.* a rabbit polyclonal antibody against phosphop44/42 MAPK and an anti-total MAP kinase antibody, were purchased from Cell Signaling Technology. a-Actin antibody was obtained from Abcam (Cambridge, UK). The IP-One ELISA assay kit from CisBio Bioassays. The fluorenylmethyloxycarbamate-(Fmoc-) protected amino-acids, Rink amide AM resin (with Nle), N,N’-diisopropylcarbodiimide (DIC) and ethyl cyanohydroxyiminoacetate (Oxyma) were purchased from Chem-Impex (Wood Dale, IL, USA). Trifluoroacetic acid (TFA), methanol (MeOH), acetonitrile (ACN), diethyl ether, N,N-dimethylformamide (DMF), piperidine, dichloromethane (DCM), and cell mask were obtained from Fisher Scientific (Nepean, ON, Canada). YM254890 was purchased from Cedarlane (Burlington, ON, Canada). All other chemicals, including 99. 9% ^2^H_2_O, were from Sigma-Aldrich (Mississauga, ON, Canada). 98% DPC-d_38_ was obtained from Cambridge Isotope Laboratories, Inc. (Andover, MA, USA), and [(2,2,3,3-tetradeuterio-3-(trimethylsilanyl)]propionic acid (TSP) from MSD Isotopes (Montreal, QC, Canada).

### Synthesis of hUT-derived pepducins

hUT-derived pepducins were synthesized manually using a standard solid phase peptide synthesis approach with Fmoc chemistry. Coupling efficiency was monitored with the qualitative ninhydrin test and a 3-equivalent excess of the protected amino acids, based on the original substitution of the resin (0.50 mmol.g^-1^), was used in most cases. Coupling of protected amino acids were mediated by DIC (1.5 eq) and Oxyma (1.5 eq) in DCM for 1 h. Fmoc removal was achieved with 20% piperidine in DMF for 20 min. All peptides were cleaved from the resin support with simultaneous side chain deprotection by treatment with TFA containing 1,2-ethanedithiol (2.5%), water (2.5%) and triisopropylsilane (1%) for 1.5 h at room temperature. The diethyl ether-precipitated crude peptides were solubilized in 70% acetic acid (1 mg/mL) and lyophilized prior to purification. Each crude peptide was purified using a preparative reversed-phase HPLC (RP-HPLC) protocol using a linear gradient from eluent A to B with 1% B per 2 min increments (Eluent A = H_2_O, 0.1% TFA, Eluent B = 80% CH_3_CN/20% H_2_O, 0.1% TFA). Homogeneity of purified fractions was assessed by RP-HPLC (eluent system: A = H_2_O (0.1% TFA) and B = 100% CH3CN with a gradient slope of 1% B/min, at flow rate of 1 mL/min on a Vydac C_18_ column) and MALDI-TOF mass spectrometry in linear mode using a-cyanohydroxycinnamic acid as matrix. Fractions presenting both the correct mass, as evaluated by MALDI-TOF mass spectrometry, and a purity higher than 98%, as confirmed by analytical RP-HPLC, were pooled, lyophilized and stored as a powder until use (**Table 1**). The synthesis of human urotensin II (hUII) was carried out as reported earlier (17,18)

### Nuclear magnetic resonance

Samples for NMR spectroscopy were prepared by dissolving the appropriate amount of peptide in 0.54 mL of ^1^H_2_O (pH 5.5), 0.06 ml of ^2^H_2_O to obtain a final DPC-d_38_ concentration of 2 mM and 200 mM. NMR spectra were recorded on a Varian INOVA 700 MHz spectrometer equipped with a z-gradient 5 mm triple-resonance probe head. All spectra were recorded at a temperature of 25°C and were calibrated relative to TSP (0.00 ppm) as internal standard. One-dimensional (1D) NMR spectra were recorded in the Fourier mode with quadrature detection. The water signal was suppressed by gradient echo (62). Two dimensional (2D) DQF-COSY (63,64), TOCSY (65), and NOESY (66) spectra were recorded in the phase-sensitive mode as described (67). Data block sizes were 2048 addresses in t_2_ and 512 equidistant t_1_ values. Before Fourier transformation, the time domain data matrices were multiplied by shifted sin^2^ functions in both dimensions. A mixing time of 70 ms was used for the TOCSY experiments. NOESY experiments were run with mixing time of 100 ms. The qualitative and quantitative analyses of DQF-COSY, TOCSY, and NOESY spectra, were obtained using the interactive program package XEASY (68). Complete ^1^H NMR chemical shift assignments were effectively achieved for hUT-Pep2 and [Trp^1^, Leu^2^]hUT-Pep3 according to the Wüthrich procedure (69) *via* the usual systematic application of DQF-COSY, TOCSY, and NOESY experiments with the support of the XEASY software package (Supporting Information, **Tables S1 and S3**). ^3^*J*_αN_ coupling constants were measured by 1D proton and DQF-COSY spectra. Temperature coefficients of the amide proton chemical shifts were calculated from 1D ^1^H NMR and 2D TOCSY experiments performed at different temperatures in the range 25-40°C by means of linear regression. Observed NOEs are reported in **Tables S2 and S4** (Supporting Information).

### Structure Determination

The NOE-based distance restraints were obtained from NOESY spectrum of hUT-Pep2 collected with a 100 ms mixing time. The NOE cross peaks were integrated with the XEASY program and were converted into upper distance bounds using the CALIBA program incorporated into the DYANA program package (Table S2) (70). An error-tolerant target function (tf-type=3) was used to account for the peptide intrinsic flexibility. From the produced 100 conformations, 10 structures were chosen, whose interproton distances best fitted NOE derived distances, and then refined through successive steps of restrained and unrestrained energy minimization using the Discover algorithm (Accelrys, San Diego, CA) and the consistent valence force field (CVFF) (71). Molecular graphics images were produced using the UCSF Chimera package (72).

### Circular dichroism

Circular dichroism (CD) analyses were performed in a JASCO J-815 (JAPAN Spectroscopy & Chromatography Technology) spectropolarimeter coupled to a temperature controller Peltier. Samples were dissolved at 0.2 mg.mL^-1^ in either water, 0.5 mg.mL^-1^ DPC (dodecylphophocholine), 0.1 mg.mL^-1^ SDS (sodium dodecyl sulfate) or TFE (2,2,2-trifluoroethanol) aqueous solutions at concentrations of 20%, 50% and 80%. The CD spectra were recorded in the measured range of 200–250 nm using a scan speed of 50 nm.min^-1^, and the baseline was corrected by subtraction of the solvent spectrum. The temperature was set to 20°C.

### Cell culture

HEK 293 cells stably transfected with the human urotensin II receptor isoform (HEK 293-hUT) were cultured in Dulbecco’s Modified Eagle’s Medium (DMEM) supplemented with 10% of heat-inactivated foetal bovine serum (FBS), 100 units/mL of penicillin, 100 μg/mL of streptomycin and 2 mM of L-glutamine. Cells were maintained as a monolayer at 37°C in a humidified atmosphere of 5% CO_2_ and 95% air and passaged by trypsinization once cells reached 70-80% confluence.

### Cell viability

HEK 293 cells were seeded at a density of 2 × 10^4^ cells/well in 96 well-plates and cultured at 37°C for 24 h. Cells were then incubated in serum free media with hUT-derived pepducins at a concentration of 10^-5^ M. Viability was assessed with a MTS kit (CellTiter 96® AQ_ueous_, Promega, Madison, Wi, USA) and a microplate reader (MTX^TC^ Revelation, Dynex Tech., VA, USA) in order to determine the optical density related to the conversion of MTS into purple-colored water-soluble formazan. Positive control (cells treated with 10% DMSO) and negative control (non-treated cells) were included in each experiment. Results were expressed as the percentage of control (non-treated cells). The presence of lactate dehydrogenase (LDH), released in the culture medium from dead cells, was also determined using a commercial LDH kit (BioVision, Miltipas, CA, USA). Positive control (cells treated with lysis buffer) and negative control (non-treated cells) were included in each experiment. Results were expressed as the percentage of the maximum LDH release following cell lysis.

### Western blotting

CHO cells stably expressing hUT (CHO-hUT; gift from Drs H. Vaudry et C. Dubessy, University of Rouen, France) were grown in 12-well plates to 70-80% confluence and starved for 6 h in a serum-free medium. For kinetic experiments, cells were incubated for different time periods (0, 5, 10, 15, 20, 40, 60, and 90 min) with a fixed concentration of hUT-Pep2 or [Trp^1^, Leu^2^]hUT-Pep3 (10^-6^ M), hUII (10^-7^ M), or PBS alone (basal control condition). To establish concentration-response curves, cells, seeded as previously described, were stimulated for 5 min with various concentrations of hUII or hUT-derived pepducins (10^-14^-10^-5^ M). When applicable, CHO-hUT cells were incubated with either pertussis toxin (PTX, 200 ng/mL) for 24 h, a specific EGFR inhibitor (AG1478; 5 μM) for 30 min or a PLCβ inhibitor (U73122; 1 μM) for 60 min prior their stimulation with hUII (10^-7^ M) or hUT-derived pepducins (10^-6^ M). Similar kinetic experiments were also performed in HEK 293-hUT cells in the presence or absence of a specific G_q_ inhibitor. Briefly, HEK 293-hUT cells, grown in 12-well plates to 70-80% confluence, were starved for 6 h in a serum-free medium. Cells were treated with YM254890 (100 nM) for 2 min and then incubated for different time periods (0, 5, 10, 15, and 20 min) with a fixed concentration of hUII, hUT-Pep2 or [Trp^1^, Leu^2^]hUT-Pep3 (10^-5^ M). Cells were then rinsed twice with ice-cold PBS and homogenized in 0.5 mL of lysis buffer (20 mM Tris, 150 mM NaCl, 1 mM EDTA, 1 mM EGTA, 1% Triton X-100, 2.5 mM sodium pyrophosphate, 1 mM β-glycerolphosphate, 1 mM Na_3_VO_4_, 1 μg/mL leupeptin, 1 mM PMSF, 1 mM DTT) at 4°C. Insoluble material was removed by centrifugation at 12,000 rpm for 30 min. The extract was treated with Laemmli sample buffer and heated for 5 min at 95°C. Aliquots were assayed for protein concentration (BioRad) and an equal amount of total protein was subjected to 12% SDS-PAGE, followed by transfer to a nitrocellulose membrane. The membrane was blocked and then probed with primary antibody (phospho-p44/42 MAPK) at the optimal dilution (1/2000) with gentle agitation overnight at 4°C. After washing with TTBS, membranes were then incubated with secondary antibody, conjugated with horseradish peroxidase (anti-rabbit HRP) diluted in TBST containing 5% skim milk (w/v) for 1 h at room temperature. Immunoreactive species were visualized by enhanced chemiluminescence using X-ray film and Thermo Scientific™ SuperSignal™ chemiluminescent substrates according to the manufacturer’s instructions. Films were digitized and the intensity of the immunoreactive bands determined using Quantity One software from Bio-Rad. To confirm equal protein loading in individual lanes, the membrane was stripped and reprobed with antibody against total ERK_1/2_ (diluted 1:5000).

### Bioluminescence resonance energy transfer experiments

HEK 293-hUT cells were grown in DMEM culture media supplemented with 10% foetal bovine serum, HEPES, sodium pyruvate and G418 (200 μg/mL). Passages were performed when cells reached 80% of confluency. Cells were initially plated in 96-well plates at a density of 10,000 cells/well. The following day, the medium is replaced with DMEM culture media supplemented with 2.5% foetal bovine serum (FBS), and 12 h later, cells were transiently transfected with 25 ng/well of a Gq-polycistronic, 25 ng/well of a G_12_-polycistronic BRET sensor, 2.5 ng β-arrestin 1-polycistronic BRET sensor, or 2.5 ng β-arrestin 2-polycistronic BRET sensor using Lipofectamine®2000 at a ratio of 3 to 1 (Lipofectamine/plasmid DNA) in serum-free Opti-MEM (73,74). Sixteen hours posttransfection, medium was replaced with DMEM supplemented with 5% FBS, HEPES, sodium pyruvate and G418 (200 μg/mL) and cell growth was resumed for another 16 h. Culture medium was changed with fresh medium DMEM/5% FBS (without antibiotics) 5 h later after transfection. Cells were washed with 120 μL PBS solution supplemented with 0.1% of glucose, and then incubated with 80 μL of this solution for 2 h. Then, to evaluate G_q_ and G_12_ activation, 10 μL of a 1/20 dilution (diluted before use) of coelenterazine 400A (stock at 1 mM in ethanol) in Kreb’s buffer and 10 μL of the 10x appropriate concentration of peptide were added, and the luminescence was evaluated with an Infinite® M1000 PRO. For β-arrestin 1 and 2 pathways, UII, URP, hUT-Pep2 or [Trp^1^, Leu^2^]hUT-Pep3 at various concentrations was first incubated for 15 min at RT prior adding coelenterazine (10 μL of a 1/20 dilution). Readings were performed after 5min using an Infinite® M1000 PRO. Filters were set at 410 nm and 515 nm for detecting the *Renilla* luciferase II (RlucII, donor) and green fluorescent protein 10 (GFP10, acceptor) light emission, respectively. BRET ratio was determined by calculating the ratio of the light emitted by GFP10 over the light emitted by the RlucII. BRET signals were normalized to that of hUII (10^-6^ M).

### IP_1_ assay

This assay, performed with the IP-One terbium immunoassay kits from Cisbio, was used according to the manufacturer recommendation and as previsouly described (43). Briefly, CHO-hUT cells were grown in 10-cm dishes for 24 h. The cells were then starved without serum. The next day, cells were washed once with PBS and collected in PBS containing 20 mM EDTA. For the assay, 80,000 cells per well (96-well plate) were used. Cells were treated for various time (10, 20, 40, and 60 min) at 37°C with hUII or treated with increasing concentrations of hUII, hUT-Pep2 or [Trp^1^, Leu^2^]hUT-Pep3 for 40 min at 37°C. IP_1_-*d_2_* and anti-IP_1_-cryptate were added for an additional 2 h at room temperature. Plates were read on a Flexstation 3 microplate reader.

### EGFR transactivation

HEK 293 and HEK 293-hUT cells, transiently transfected with cDNA encoding fluorescently labeled EGFR (EGFR-GFP) (2 μg), were plated onto 35-mm glass-bottomed culture dishes (MatTek Corp, Ashland, MA). Following stimulation, cells were washed once with PBS and fixed in 4% paraformaldehyde for 15 min. To study the involvement of G_q_ in EGFR internalization, cells were treated with YM254890 (100 nM) for 1h prior peptide treatments. EGFR internalization following hUII, hUT-Pep2, or [Trp^1^, Leu^2^]hUT-Pep3 stimulation (10^-6^ M) was visualized by green fluorescence using a single sequential line of excitation on a Zeiss LSM 780 Axio Observer Z1 (Carl Zeiss Microimaging). EGFR-GFP internalization was visualized using a combination of excitation (488 nm) and emission filters (499 and 520 nm).

### Proliferation assays

HEK 293-hUT cell proliferation was evaluated by flow cytometry using propidium iodide DNA staining. Cells, grown in 6-well plates to 80% confluency, were starved for 6 h in a serum-free medium and then treated with hUII, hUT-Pep2, or [Trp^1^, Leu^2^]hUT-Pep3 (10^-6^ M) for 30 minutes at RT. Next, cells were detached, fixed with cold 70% ethanol and then stored at −20°C for at least 2 h. Cells were centrifuged at 200 rpm for 10 min at 4°C and washed at least once with cold PBS. Cells were then resuspended in Triton X-100 (0.1%) supplemented with propidium iodide (500 μg/mL), and 200 μg/mL of DNAse-free RNAse A in PBS. Data, expressed as the mean ± SEM of at least three independent experiments, were acquired on a FACScan (BD Biosciences, San Jose, CA, USA) and results were analyzed with the FlowJo v10 software. The proliferation index is defined as the total number of divisions divided by the number of cells that went into division.

### Rat aortic ring contraction

Adult male Sprague-Dawley rats (Charles-River, Saint-Constant, Qc, Canada) weighing 250-300 g were housed in cages under controlled illumination (12:12 h light-dark cycle), humidity, and temperature (21-23°C) and had free access to tap water and rat chow. All experimental procedures were performed in accordance with regulations and ethical guidelines from the Canadian Council for the Care of Laboratory Animals and received approvals of the institutional animal care and use committee of the *Institut National de la Recherche Scientifique - Centre Armand-Frappier Santé Biotechnologie*. As previously described,(17,18) the thoracic aorta was cleared of surrounding tissue and then excised from the aortic arch to the diaphragm. Conjunctive tissues were next removed from the thoracic aorta and the vessels were divided into 5 rings of 4 mm. The endothelium of each aortic ring was removed by gently rubbing the vessel intimal surface. Aortic rings were then placed in a 5 mL organ bath filled with oxygenated normal Krebs-Henselheit buffer. Eighty microliters of a 2.5 M potassium chloride solution (40 mM final concentration) were used to evaluate maximal contractile responses of each vessel. In one bath, hUII (10^-6^ M) was applied as a control, and the tissue-response was expressed as the ratio with the KCl-induced contraction. Cumulative concentration-response curves to synthetic peptides were obtained by increasing the concentration of each peptide in the remaining organ chambers (10^-11^ to 10^-5^ M). The amplitude of the contraction induced by each concentration of peptide was expressed as a percentage of the KCl-induced contraction divided by the tissue-response induced by hUII. Antagonist activity of hUT-derived pepducins was evaluated by exposing first the aortic ring to hUT-Pep2, or [Trp^1^, Leu^2^]hUT-Pep3 (10^-5^ M) for 30 min, to ensure that the peptide reached equilibrium and that no agonist effect is observed, and then adding cumulative concentration of hUII or URP (10^-12^-10^-6^ M). The median effective concentrations (EC_50_) are expressed as the mean ± S.E.M., and the *n* values, representing the total number of animals from which the vessels were isolated, varied from 5-8 animals.

### Rat neonatal fibroblast proliferation

Neonatal rat fibroblasts, endogenously expressing UT (75), were obtained from 1-3 days old Sprague-Dawley pups as previously reported (76). Briefly, hearts obtained from neonatal rats were enzyme-digested and after centrifugation, cells were pre-plated to separate cardiomyocytes from fibroblasts. As reported, the adherent nonmyocyte fractions correspond predominantly to fibroblasts while the supernatants largely contain the cardiomyocytes. After several washes, attached fibroblasts were trypsinized and then cultured at 37°C in 96-well plate (6,000 cells per well) in low glucose-DMEM supplemented with 7% FBS, 100 units/mL of penicillin, and 100 μg/mL of streptomycin. After 48 h, cells were washed twice with low glucose-DMEM, and starved for 12 h in this medium. Cells were then treated for 24 h with hUII, hUT-Pep2, or [Trp^1^, Leu^2^]hUT-Pep3 at a concentration of 10^-6^ M. Prior to staining, cells were fixed with 4% paraformaldehyde for 15 min at RT, permeabilized with 0.2% Triton X-100 in PBS for 5 min, and then blocked with 10% horse serum in PBS for 1 h at RT. Cells are incubated with primary anti-smooth muscle a-actin antibody (1/500) in 10% horse serum/PBS overnight at 4°C and later with secondary anti-rabbit Alexa Fluor 488 (1/1000) for 1 h at RT. Cell staining was performed using cell mask (1 μg/mL) in PBS for 30 min at room temperature and with Hoechst dye in PBS (1 μg/mL) for 10 min. Analysis was performed on an Opera Phoenix High Content Imaging System with a 20x water objective.

## Data availability

The ensemble of hUT-Pep2 NMR-derived structures has been deposited in the Protein Data Bank (PDB) with the following code: 6HVK. All remaining data are contained within the article.

## Statistical analysis

All experiments were performed at least in triplicate. Data, expressed as mean ± S.E.M, were analyzed with the Prism Software (Graphpad Software, San Diego, CA, USA). Sigmoidal concentration-response curves fitted with variable slope were used to determine EC_50_. Statistical comparisons were analyzed by the Student’s t-test, and differences were considered significant where \**P* < 0.05, \*\**P* < 0.01 or \*\*\**P* < 0.001.

## Acknowledgements

This work was supported by a grant from the Heart and Stroke Foundation of Canada (G-15-0008938) and the Canadian Institutes of Health Research (PJT-159687) to T.E.H. and J.C.T, a grant from the Canadian Institutes of Health Research to D.C. (MOP-142184) and a grant from the Natural Sciences and Engineering Research Council of Canada (RGPIN-2015-04848) to D.C. R.D.M. was supported by studentships from the McGill-CIHR Drug Development Training Program and the McGill Faculty of Medicine and Health Sciences. H.N. was supported by a fellowship from the *Fonds de Recherche en Santé du Québec*. J.C.C.D was supported by the Barbara & Edward Victor award granted by McGill Faculty of Medicine.

## Conflict of interest

The authors declare that they have no conflicts of interest.

## Abbreviations

7TMR: 7-transmembrane receptor
BRET: bioluminescence resonance energy transfer
BOP: (benzotriazol-1-yloxy)tris(dimethylamino) phosphonium hexafluorophosphate
CHO cells: Chinese hamster ovary cells
DCM: dichloromethane
DIEA: N,N-diisopropylethylamine
DMF: dimethylformamide
DPC: dodecylphosphocholine
DQF-COSY: double quantum filtered correlated spectroscopy
Fmoc: fluorenylmethyloxycarbamate
GPCR: G protein-coupled receptor
HEK 293 cells: human embryonic kidney 293 cells
MALDI-TOF: matrix-assisted laser desorption/ionization-time of flight
NMR: nuclear magnetic resonance
NOESY: nuclear Overhauser enhancement spectroscopy
RP-HPLC: reverse phase-high performance liquid chromatography
TOCSY: total correlated spectroscopy
TSP: 3-(trimethylsilanyl)propionic acid
UII: urotensin II
URP: urotensin II-related peptide
UT: urotensin II receptor

